# Melanism in a wild sifaka population: Darker where cold and fragmented

**DOI:** 10.1101/2022.02.02.478891

**Authors:** Elizabeth Tapanes, Jason M. Kamilar, Maanas A. Nukala, Mitchell T. Irwin, Brenda J. Bradley

**Affiliations:** Center for the Advanced Study of Human Paleobiology, Department of Anthropology, The George Washington University; Graduate Program in Organismic and Evolutionary Biology University of Massachusetts Amherst; Department of Anthropology, University of Massachusetts Amherst; Department of Anthropology, Northern Illinois University

**Keywords:** ecological niche modeling, Indriidae, coloration, lemurs, trait evolution, conservation

## Abstract

Pigmentation is one of the most striking examples of diversity in the natural world. Mainly, pelage (hair/fur) pigmentation provides a substrate for selection (i.e., crypsis, signaling, thermoregulation) and is capable of rapid change. Thus, this trait may be the one potential early signal of adaptation (or maladaptation) in wild primate populations. However, most of our hypotheses on the forces responsible for primate pelage pigmentation are based solely on macro-evolutionary studies. Here, we characterize pelage color and pattern variation within a population of wild primates, diademed sifakas (*Propithecus diadema*), exhibiting striking diversity in coloration (melanic to tri-colored). Our approach jointly assesses climate and pelage variation across the region. We score pelage using a semi-quantitative methodology. We then test if pelage variation is associated with climatic or demographic factors (i.e., sex-class, age-class) across the Tsinjoarivo forest, Madagascar. We find darker bodies and less complex faces occur in colder and more fragmented forests. We explore three hypotheses that may explain this phenotypic pattern: isolation by distance, an environmental gradient, or unique local adaptation. Importantly, each scenario signals the need for enhanced conservation of diademed sifakas in the Tsinjoarivo forest. More studies on primate pigmentation in wild populations will be needed to contextualize if this pattern is exceptional or typical. It is likely that in other primate populations pigmentation may also foretell of adaptation or environmental mismatch.

## Introduction

Variation in pigmentation of pelage (hair, fur) exemplifies some of the most striking observable diversity in mammals. Primates, in particular, exhibit some of the most remarkable diversity in pelage pigmentation, displaying a high degree of inter- and intraspecific variation (Bradley & Mundy, 2008; Caro et al., 2017). Pelage color patterns span black and white variegated patterns (e.g., black-and-white ruffed lemur (*Varecia variegata*)), orange hues (e.g., orangutans (*Pongo* spp.)), stripes (e.g., ring-tailed lemurs (*Lemur catta*)), and intricate facial patterns (e.g., guenons (*Cercopithecus* spp.)) (Bradley & Mundy, 2008). Current hypotheses argue that pelage pigmentation aids with crypsis, signaling, and/or thermoregulation (Caro et al., 2021; Cuthill et al., 2017; Tapanes et al., 2021). However, even against a backdrop of selection, neutral factors such as drift and restricted gene flow can influence pigmentation patterns (Runemark et al., 2010). Nevertheless, darker species tend to co-occur in warm, humid forests, and species with more complex color patterns tend to live in larger groups or exhibit enhanced color vision (Allen et al., 2014; Bell et al., 2021; Santana et al., 2012; 2013). These patterns may implicate camouflage and crypsis, respectively. Additionally, many species exhibit marked ontogenetic coat changes (Caro & Mallarino, 2020). However, most of what we know about forces potentially driving variation in primate hair pigmentation derive from macro-evolutionary studies across large portions of the order (e.g., all African monkeys and apes) (Caro et al., 2021; Santana et al., 2013).

What drives variation on a micro-evolutionary scale is not always identical to what drives macroevolutionary variation. For example, human hair color variation likely evolved due to neutral forces such as drift—but, in general, variation across the primate order exhibits evidence of potential adaptive evolution (Caro et al., 2021; Jablonski & Chaplin, 2017; Santana et al., 2013; Tapanes et al., 2021). In both neutral and non-neutral evolution, hair pigmentation is often capable of rapid change. When adaptive, variation in hair within a population can mark the difference in an individual’s survival or demise. For example, snowshoe hare predation increases when a mismatch exists between coat color and snow cover (Zimova et al., 2016). So, in response to milder winters in the Pacific Northwest, hare populations are now rapidly evolving brown winter coats (Jones et al., 2018). In primates, mantled howler monkey (*Alouatta palliata*) pelage color recently experienced rapid change—likely due to an environmental shift (Galván et al., 2019). Whether or not this will lead to environmental mismatch or rapid adaptive evolution is yet to be seen. Mainly, this is due to a lack of data on the forces and mechanisms that drive hair variation within primate populations. Thus, studying population-level pelage color broadly relates to understanding wild primate adaptation (or maladaptation).

Indriidae, a lemur family, is an ecologically diverse radiation of primates in Madagascar that exhibits spectacular pelage diversity. Specifically, the coloration of sifakas (*Propithecus* spp.) has been the subject of speculation for decades because of high diversity in pelage color patterns with variable degrees of black coloration, both among species and within populations (Mittermeier et al., 2010). Naturalists have documented melanic (black) morphs since the mid-1900s in various sifaka populations—across species that are not typically melanic. Early naturalists and recent empirical evidence show dark morphs co-occur in colder forests (King et al., 2014; Pastorini et al., 2001; Petter & Peyrieras, 1972; Tapanes et al., 2021). This pattern may implicate selection for enhanced thermoregulation (Bogert’s rule) because dark coat colors may aid in absorbing heat from the sun’s rays (Bogert, 1949; Delhey et al., 2019). However, some of these populations have already gone locally extinct due to intense deforestation in Madagascar (Tattersall, 1986).

Nevertheless, there is a unique opportunity to study hair color variation in a population of diademed sifakas (*Propithecus diadema*) living in the unprotected forest of Tsinjoarivo, Madagascar (19°41’S, 47°48’E). Here, sifakas exhibit an extraordinary degree of color variation and facial patterning complexity (Figure 1) (Mittermeier et al., 2010), and have been under study for 20+ years (Irwin, 2006; Irwin et al., 2019). Throughout most of the species’ range, diademed sifakas exhibit a tri-colored orange, black, and white pelage. At Tsinjoarivo, unique coloration patterns include the classic tri-colored morphotype, completely melanic individuals, partly melanic individuals (e.g., melanic from the dorsal torso upwards), and variations in between (Mittermeier et al., 2010). Historically, geneticists considered placing Tsinjoarivo sifakas in a distinct taxon (Rumpler et al., 2011) because their unique pelage coloration patterns are not typical of the rest of the species’ range (Groves, 2001). However, Tsinjoarivo sifakas are currently subsumed into diademed sifakas and recognized as a non-distinct genetic population-based on mitochondrial DNA (Mayor et al., 2004). Sifakas at Tsinjoarivo live in both continuous and fragmented forest. Tsinjoarivo is also typified by a significant elevational gradient–highest in the west (≈1600m) and lowest in the east (≈1350m). Previous climate recordings from inside the forest indicate that more rain falls in the eastern-most block of continuous forest on average (≈2600mm) in comparison with fragmented forest patches in the west (≈2000mm) (Irwin, 2006). Thus, Tsinjoarivo sifakas represent an ideal case to test what potential forces (e.g., climatic, demographic) underly primate coloration variation within a wild primate population.

**Figure 1.**
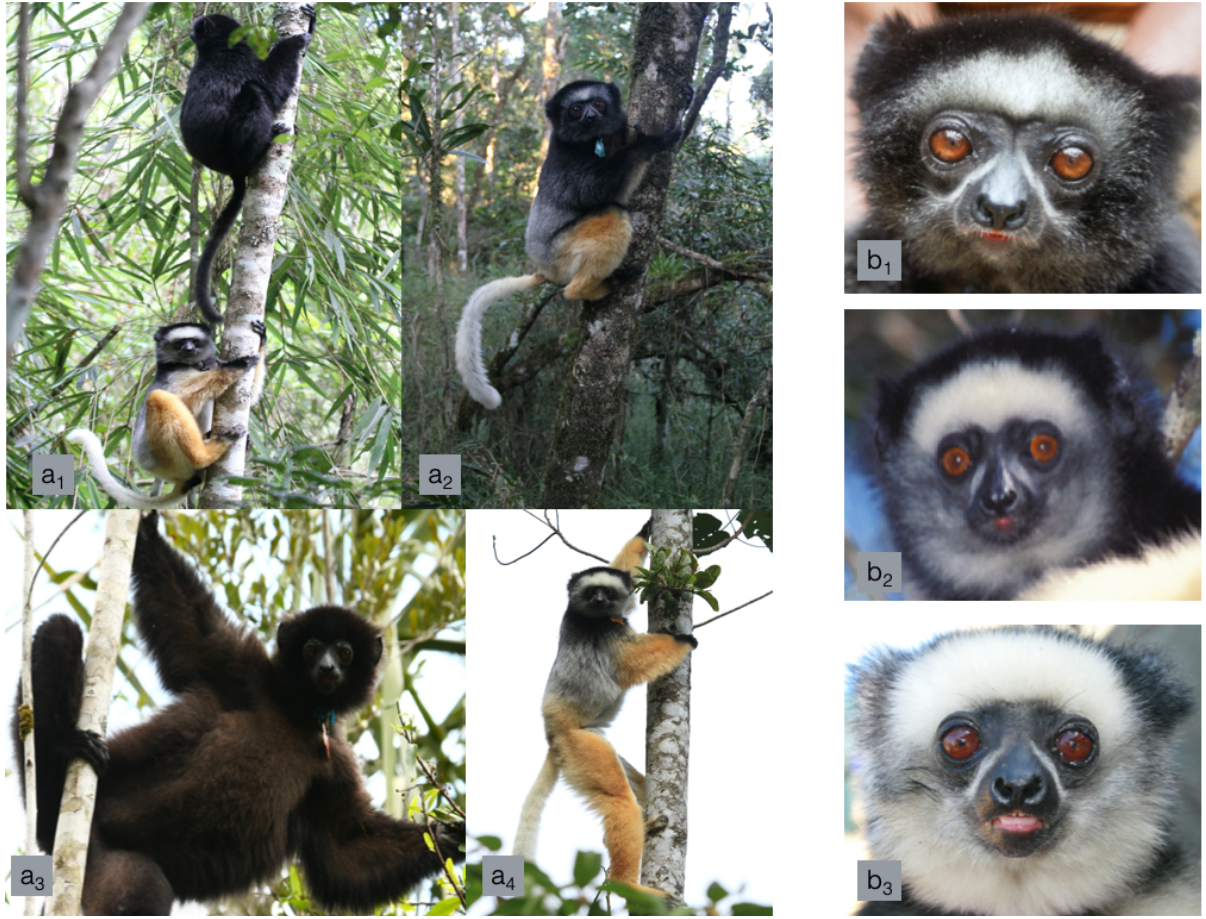
Pelage color variation in diademed sifakas of Tsinjoarivo, Madagascar (a) across the anterior body and (b) across the head

In this study, we first examined climatic variation across the Tsinjoarivo forest. We were interested in determining whether variation in climatic regions, or lack thereof, could provide insight into pelage variation. We hypothesized we would find significant climate differences, based on previously recorded data at Tsinjoarivo that has not been empirically tested (Irwin, 2006). We then analyzed photographs of wild sifakas living in Tsinjoarivo over the past ≈20 years to test how climate and/or demography co-vary with whole-body pelage pigmentation and pattern complexity. We predicted that pelage morphs would be distributed randomly throughout the forest regardless of climate, consistent with neutral evolution. Alternatively, we predicted pelage morphs may be distributed in a non-random pattern. The latter would implicate either selection pressure or restricted gene flow.

## Methods

### Climate Modelling

#### Study site

At Tsinjoarivo, we sampled fragmented (towards the west) and continuous forest (towards the east) through three base camps previously set up at Tsinjoarivo. From west to east, the first is Mahatsinjo camp (i.e., FRAG1-7, 19°40.94’S, 47°45.46’E) in the west, followed by Ankadivory camp (i.e., CONT4-5, 19°42.98’S, 47°49.29’E), and lastly by Vatateza camp (i.e., CONT1-3, 19°43.25’S, 47°51.41’E) (Figure 2). In general, the eastern-most continuous section of the forest has a higher canopy, fewer but larger trees, a higher basal area per hectare, and a higher quality forest compared to the western-most fragmented forest (Irwin, 2006; Irwin et al., 2019).

**Figure 2.**
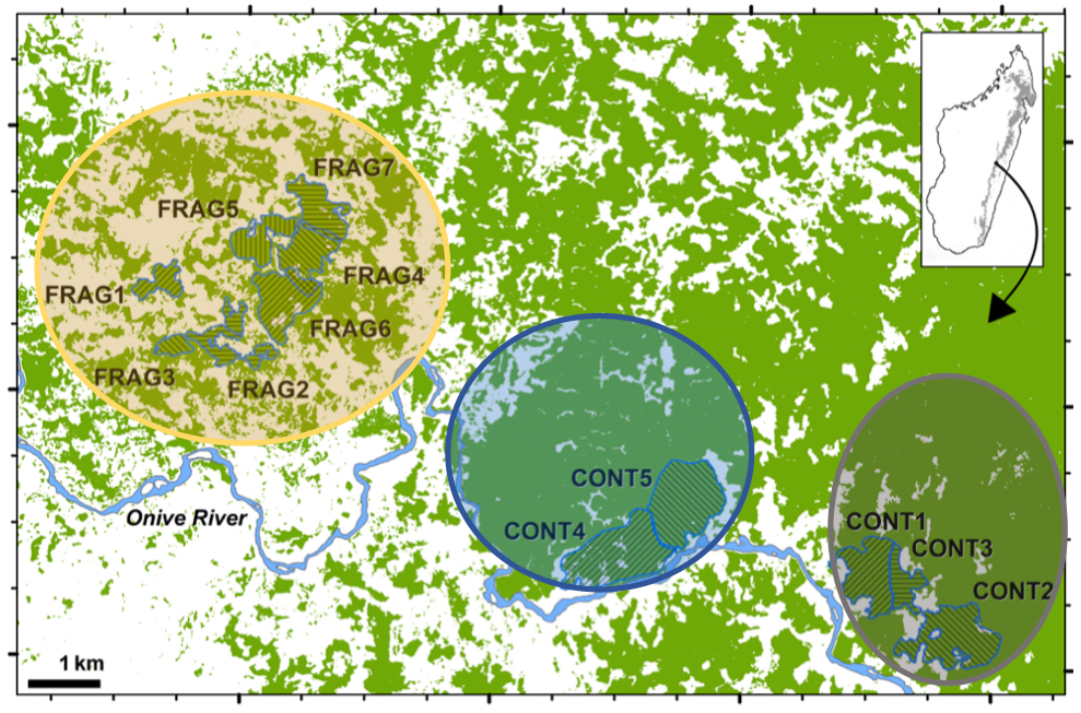
A detailed map of the spatial composition of all groups in Tsinjoarivo, Madagascar across three base camps. The yellow circle encompasses Mahatsinjo, the blue circle encompasses Ankadivory, and the grey circle encompasses Vatateza.

#### Analyses

We collected climate data using presence-only occurrence points for 13 sifaka groups across Tsinjoarivo, including seven groups from the fragmented forest and five groups from the continuous forest, using WorldClim version 1 (Hijmans et al., 2005) at 30-sec (ca. 1km) spatial resolution. WorldClim is widely used in biogeography studies (Fuchs et al., 2018) and has been used effectively in Madagascar despite the lower number of weather stations (Pearson, 2015; Vieilledent et al., 2013). We confirmed three extremely close (≈3km) weather stations to Tsinjoarivo and an additional six only ≈30-40km in either direction (NE, SE, NW, SW). We clipped the data to the species range but extended it out to the coast (using QGIS v3.10.1) because some diademed sifaka populations currently live in forest remnants closer to the coast (i.e., Betampona). Additionally, multiple Indriidae species likely occurred outside their current ranges as little as 20-50 years ago due to rapid deforestation in Madagascar, especially the eastern lowlands (Jungers et al., 1995).

We ran a correlation matrix of all 19 WorldClim bioclimatic variables (measuring various aspects of rainfall and temperature) extracted across habitats in Tsinjoarivo (from the 13 groups) and removed variables that exhibited correlation coefficients > 0.80. The resulting dataset included six bioclimatic variables (SI: Table 1). We generated Ecological Niche Models (ENMs) using Maxent v3.4.1 (Phillips, Dudik, & Schapire, n.d.) with default settings. We estimated model performance using the area under the receiver operating characteristic curve (AUC), and took models above 0.70 to indicate good performance (Swets & Swets, 1988). ENMs can be used to assess the predicted distribution of entire species (Elith & Leathwick, 2009; Elith et al., 2011). However, recent studies indicate that ENMs on populations provide data on population differentiation because subtle environmental differences can adequately reflect local variation (Gonzalez et al., 2011; Soto-Centeno et al., 2013).

**Table 1.**
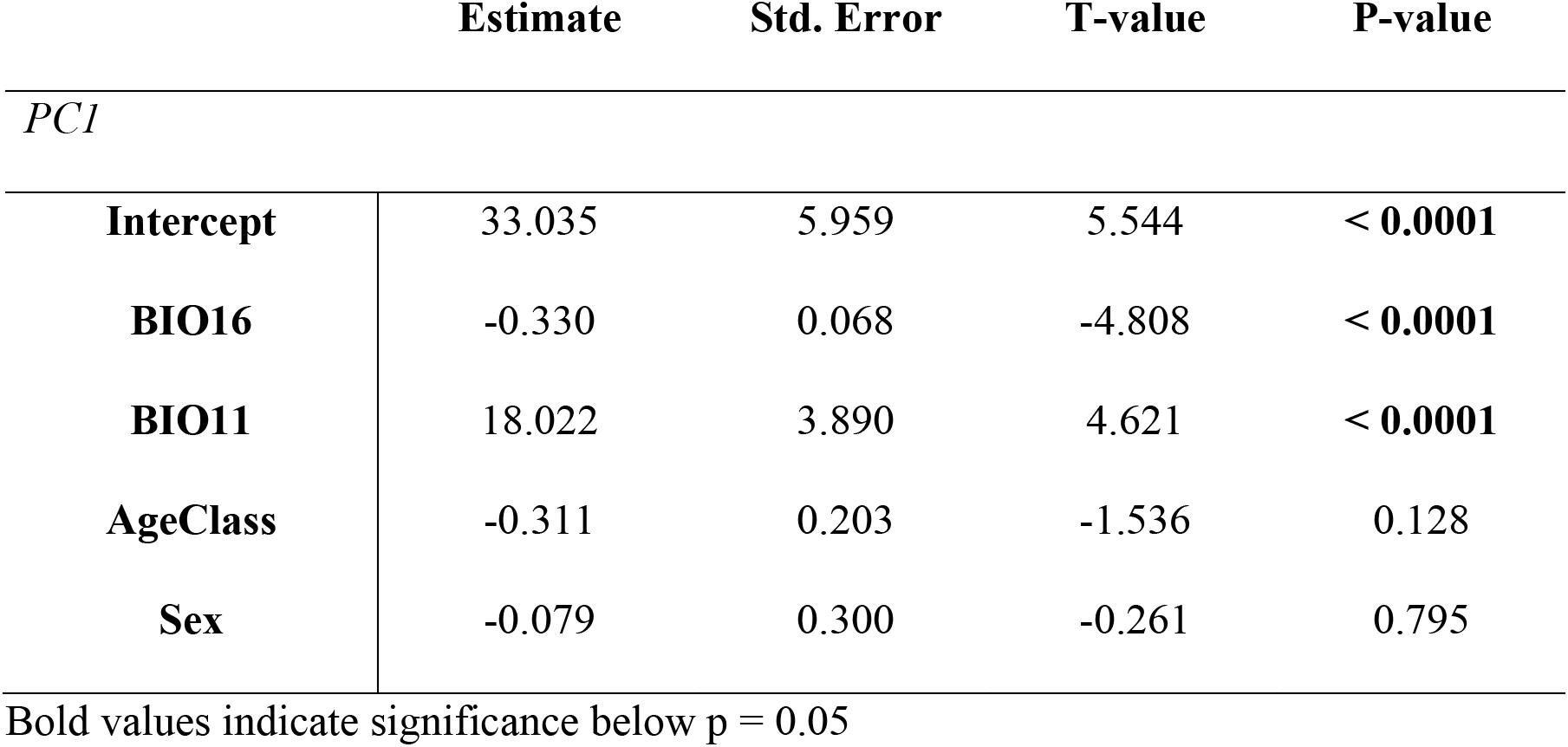
General linear model (GLM) predicting the PC1 values of the pigmentation PCA while including all independent variables.

After generating ENMs, we used an identity test to assess the similarity between models developed from fragmented vs. continuous habitat using the R package ‘ENMTools’ (Fuchs et al., 2018; Warren et al., 2010). We did not assess the differences between each ‘region’ because we lacked statistical power to divide an already small sample. The test took locality data from continuous and fragmented habitat groups and randomly assigned localities to “pseudo” habitat pairs (also known as “pseudo” species pairs in macro-scale studies). The test then took known locality points and randomized their identities to produce a new dataset composed of the same number of localities as the original input data. In total, we generated 99 pseudo habitat pairs and then compared pseudo-pairs to the real habitat data. We supplemented this with estimates of predicted similarity between both habitats using the niche overlap function in ENMtools based on *Schoener’s D* (Schoener, 1968) and *Hellinger’s I* (both range from 0 to 1) (Warren et al., 2010). A value of one indicates complete niche overlap, and zero indicates no niche overlap. Similarly, we generated 99 pseudo habitat pairs for *D* and *I* and compared observed values to the random distributions. If the observed values fell below the randomized values, no significant niche or climate overlap existed.

To further assess the Maxent models’ success, we performed a binomial omission test under a minimum training presence, at an alpha of 0.025. We followed a four-fold cross-validation approach. Due to our dataset’s small size, we withheld approximately 40% of occurrences in each habitat type as test data (e.g., 2 of 5 of continuous occurrences, 3 of 7 of fragmented). We randomly shuffled occurrences into test and train splits for each fold. Lastly, in our pelage analysis below, we used the two climate variables with the highest permutation of importance (SI: Table 2).

**Table 2.**
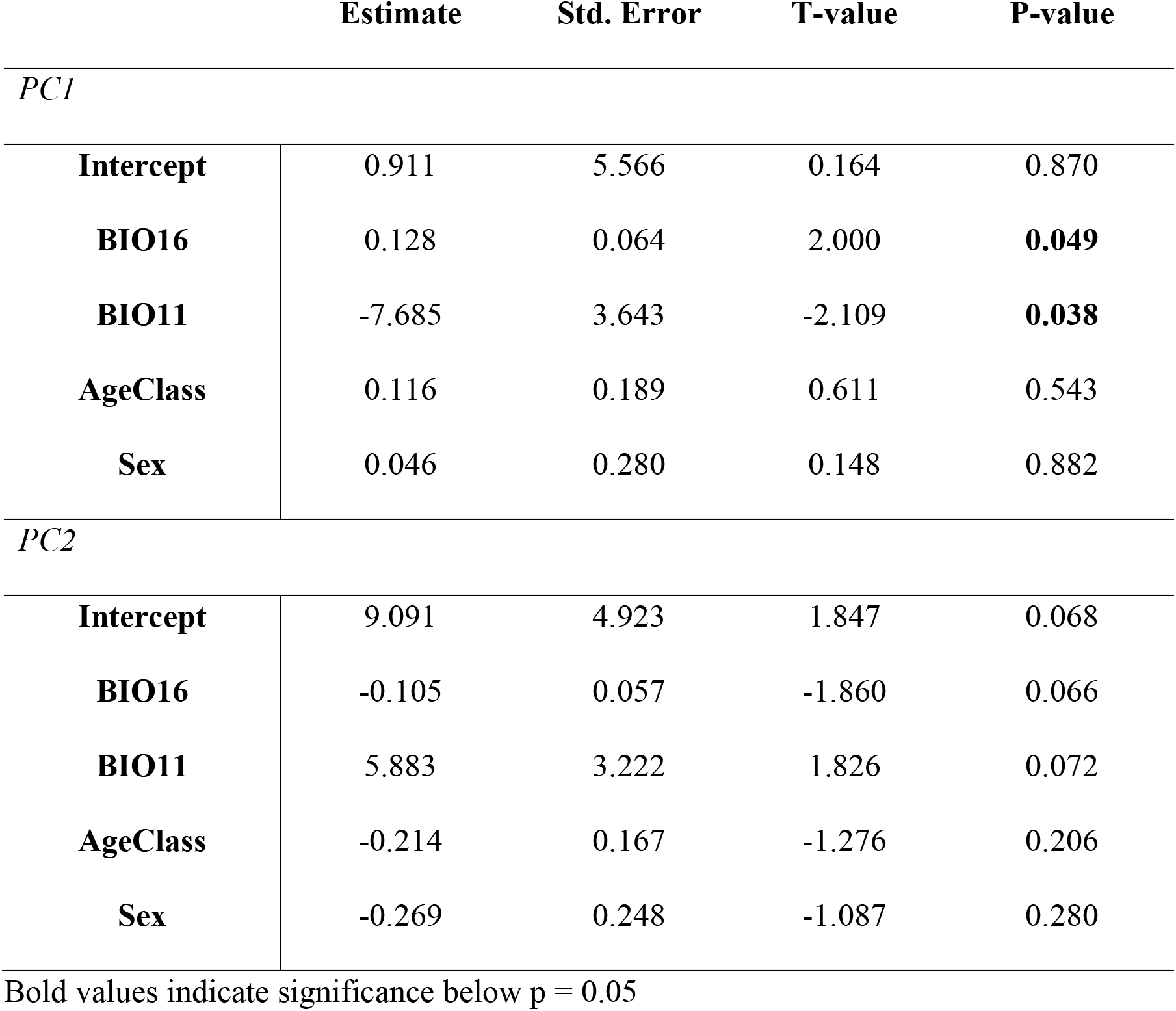
General linear model (GLM) summary predicting PC1 and PC2 of the color patterning PCA using all independent variables.

### Pelage variation

#### Sampling and Photo Collection

In this study, we examined hair color variation across Tsinjoarivo sifakas. We habituated groups before capturing and then captured animals at close range using Pneu-dart 9mm disposable non-barbed darts. A trained professional performed the immobilization with tiletamine/zolazepam (Telazol). Medical examinations followed a standard protocol established by the Prosimian Biomedical Survey Project (PBSP) Team and used on 750+ lemurs, over 16 sites, and 35 species since 2000 (Dutton et al., 2003; Irwin et al., 2010).

We took ≈20-30 digital photographs and/or slides of each sifaka during captures using an existing standard protocol (Irwin et al., 2010). Our sampling thus spanned ≈20 years’ worth of photos across different cameras and lighting conditions (N =87; SI: Table 3). We did not remove any phenotypic outliers because that would require removing the most exciting variants (i.e., all melanic) in the sample and thus negating the purpose of the study.

#### Ethical Note

We performed the research according to Madagascar’s ethical and legal requirements and the Code of Best Practices for Field Primatology. The study was ethically reviewed and approved by the IACUC (Institutional Animal Care and Use Committee) of Northern Illinois University (LA12-0011), University of Queensland, McGill University, and Stony Brook University. A team captured sifakas for ongoing parallel research, including population monitoring, sample collection, and conservation efforts—and not to study their hair variation. Therefore, our photographic sampling was opportunistic.

#### Semi-Quantitative Scoring

We developed a systematic method of scoring pelage variation semi-quantitatively (Figure 3) based on similar pelage scoring systems from previous studies in other primates (Caro et al., 2021; Rakotonirina et al., 2017; Santana et al., 2012; 2013). Two individuals scored sifaka photographs, and we verified intra- and interscorer reliability above 95%. Photos were blind coded. One scorer was unaware of hypotheses behind the study and locality of individual sifakas. The second scorer was unaware of the locality of the sifaka photos (but aware of hypotheses). We scored 18 Tsinjoarivo sifaka body regions on coloration, complexity, and contrast across a large sample (N = 87; SI: Table 4).

**Figure 3.**
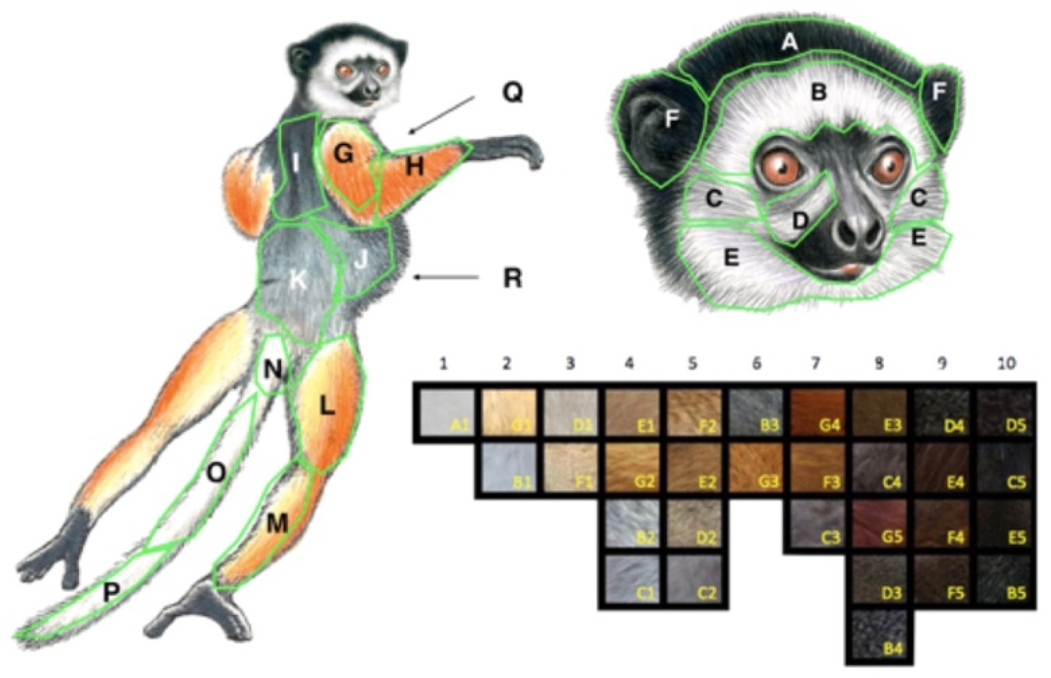
Scoring schematic use to characterize pelage variation in this study. The scoring chart was adapted from a previous study (Caro et al., 2017). Sifaka illustrations modified with permission, copyright 2013 Stephen D. Nash.

For every sifaka, we assigned each region a score (1-10) and a shade (31 possible; A1-G5) from 10-15 photos. For example, a completely orange lower forelimb (region H) scored a “6” in coloration and “G3” as the shade (Figure 3). We used numerical scores to calculate mean pigmentation scores, and letter scores (e.g., A1) to calculate complexity and contrast scores. We defined pattern complexity as the total number of shades (31 possible) across each central body region (e.g., head), and contrast as the difference in the most extreme shades of each significant body region (on a scale of 1-10). We estimated body coloration (brightness/darkness) as the mean numerical score (1-10) of all shades per area, per individual (SI: Table 4). We measured complexity and contrast using subsumed scores from the head, anterior body, and ventral-torso scores. We measured pigmentation from subsumed variables from the head, dorsal-torso, hindlimb, forelimb, ventral-torso, and tail regions. We measured countershading as the ratio between the ventral and dorsal pigmentation scores. For any individuals that were captured twice (across distinct years), we computed their mean pigmentation score (1-10) to assess brightness/overall color (i.e., darkness). Color scores did not vary over time. In body regions where more than one color/shade was evident in the coat/body region, we assigned scores based on the predominant color (Caro et al., 2017).

We note that despite criticisms that the human visual system distorts animal colors (Stevens et al., 2009), the human visual system may be good at detecting animal color variation in the general visible spectrum. Specifically, recent work shows that color extracted from images (using sets of ≈12, as we did) can be reliable—this may be particularly true for estimates, such as these, of dark or light coloration (Higham, 2021; Laitly et al., 2021).

#### Analyses

We conducted two principal components analyses (PCA) on numerical pelage variables using R’s prcomp function (R Core Team, 2018). We removed pelage variables with highly correlated vectors (i.e., overlapped in the biplot) to reduce the dimensionality of the data and meet parametric testing assumptions. The first PCA included body pigmentation (brightness/darkness) scores, and the second PCA included pattern complexity (including contrast and countershading). Henceforth, we will refer to the first as the ‘brightness PCA’ and the latter as the ‘pattern PCA.’ We used transformed PC scores (eigenvalues > 1) as dependent variables (Kaiser, 1960) in general linear regression models. We positivized all PC values by adding a constant (the absolute value of the lowest PC score) to each value.

We ran general linear models that included several predictor variables. To assess impacts from climate/ecology, we used the two climate variables that contribute to climate differences across Tsinjoarivo (along with age-class and sex-class) as our independent variables. We used the dredge function in the MuMIn package (Barton, 2019) for model selection based on Akaike’s Information Criterion with modification for small sample size (AICc). AICc scores provided us with the likelihood of a model given a dataset while minimizing model complexity. Models within two AICc values of the “best model” were deemed equally good (Burnham & Anderson, 2003; Wheeler et al., 2011). We calculated each model’s weight as a supplement to determining the best fit model, which was particularly useful when multiple models had similar AICc scores. Then, we assessed each predictor variable’s importance in our models based on their sum of AICc weights. We used G*Power (Faul et al., 2009) to determine our linear models’ statistical power. Given our sample size, the number of predictor variables, and the use of a two-tailed general linear regression, we estimated we could detect a small effect size (.29) with 85% power.

### Data Availability Statement

The datasets generated and analyzed for the current study are available from the corresponding author on reasonable request.

## Results

### Climate Modelling

Models had a mean AUC value above 0.98—which indicated high performance. Our models showed a significant climate dissimilarity between fragmented forest groups in the west and the east’s continuous forest groups (Figure 4). This gradient is potentially formed because continuous forest sits at a lower elevation than fragmented forest. Our observed values of *Shoener’s D* and *Hellinger’s I* were outside the range of the random distribution of values (SI: Figure 1) and statistically distinct (identity test, p = 0.01) (SI: Table 5). Additionally, the binomial test of omission was significant across continuous and fragmented forests across all four-folds (p < 0.0001) (SI: Table 6).

**Figure 4.**
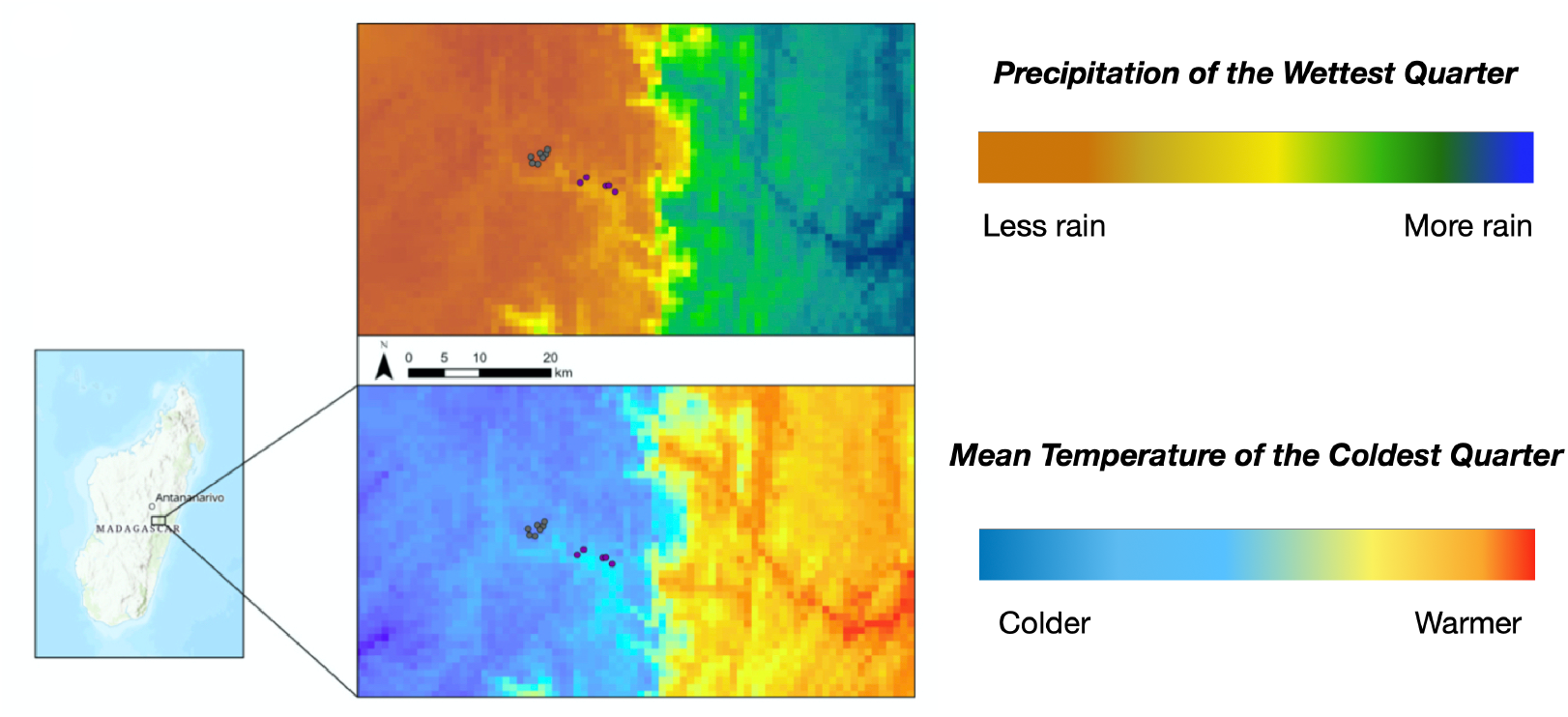
A west-east gradient that is formed in Tsinjoarivo, based on ‘Precipitation of the Wettest Quarter’ and ‘Mean Temperature of the Coldest Quarter’

Several variables distinguished fragmented and continuous forest habitat models. The highest permutation importance to the fragmented forest habitat model included precipitation of the wettest quarter (bio16) at ≈38%, but for the continuous forest, it was mean temperature of the coldest quarter (bio11) at 56%. Temperature annual range (bio07) was the second-highest contributor to both models’ permutations at 33% (SI: Table 4). These variables indicate a west-east gradient in precipitation and temperature across Tsinjoarivo (Figure 4).

### Pelage variation

Regarding the ‘brightness PCA,’ we found that the first PC (eigenvalue > 1) explained ≈62% of the variation in the dataset (Figure 5). Loadings on PC1 included forelimb coloration (.29), dorsal torso coloration (.25), ventral torso coloration (.24), and tail coloration (.20). We found a significant association between PC1 and both climate variables (precipitation of the wettest quarter (BIO16) and temperature of the coldest quarter (BIO11)) (Table 1). From this point forward, we refer to these climate variables as ‘precipitation’ and ‘temperature,’ respectively. We note that PC1 increases as temperature increases (i.e., pelage gets lighter) (Table 1, Figure 5). Specifically, this ‘brightness PCA’ illustrates that darker individuals in our sample most often lived in colder forest patches that exhibited less precipitation (Figure 5). In the best-fit models for PC1, according to AICc scores, the strongest predictors were climate variables and age class (SI: Table 7). However, our general linear model detected no significant differences across sex-class or age-class (Table 1). Additionally, both climate variables contributed ≈50% and ≈75% more to the model than either age-class or sex-class, respectively (SI: Table 8).

**Figure 5.**
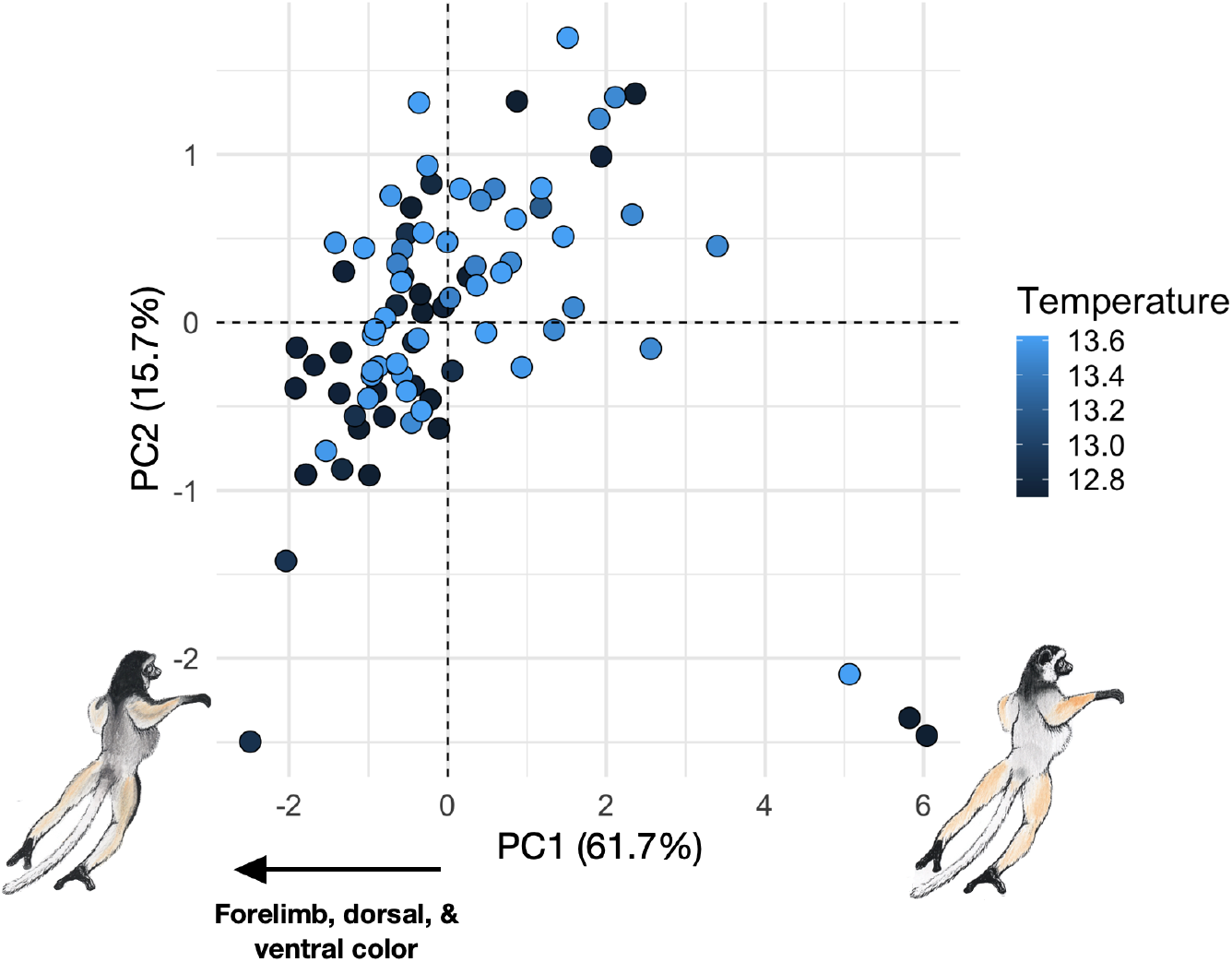
Plot of PCA scores representing overall pelage coloration of Tsinjoarivo sifakas across the climate gradient (shown with temperature). Brightness increases with increasing PCA values (and darker sifakas are represented by lower PC values).

For the ‘pattern PCA’ (to assess complexity), the first two PCs (eigenvalues > 1) explained ≈60% of the variation in the dataset (Figure 6). Highest loadings on PC1 included head color complexity (.39), head color contrast (.37), ventral torso color contrast (.24), ventral torso color complexity (.0), and countershading (.0). Specifically, we detected a statistically significant effect of temperature and precipitation on PC1 (Table 2), with temperature being negatively related to PC1 (Table 2). This indicates PC1 would be expected to increase (i.e., complexity increase) as temperatures increase, as indicated by the PCA (Figure 6).

**Figure 6.**
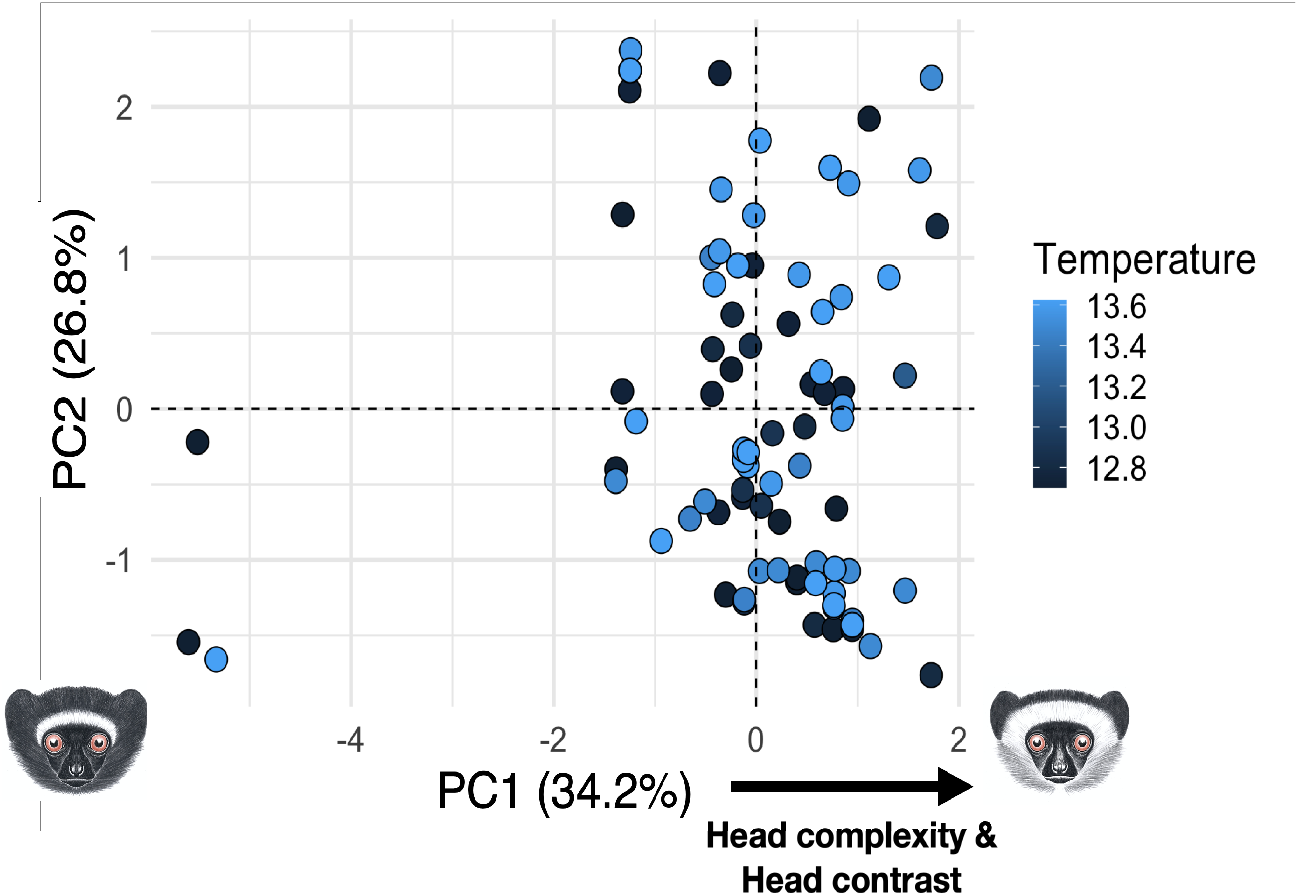
Plot of PCA scores representing the pelage complexity of Tsinjoarivo sifakas across the climate gradient. Less complex pelage is represented by lower PC values.

Furthermore, we did not detect a significant association between PC1 and age or sex class. According to AICc scores, only the two climate variables contributed to the best fit model for PC1 (SI: Table 9). Climate variables contributed ≈30% and 40% more to PC1 than either sex-class or age-class, respectively (SI: Table 10). Loadings on PC2 include countershading (.45), ventral torso color complexity (.45), ventral torso color contrast (.08), head color contrast (.03), and head color complexity (0). We did not detect statistically significant effect of any predictor on PC2. Climate variables alone accounted for the model with the second-best fit for patterning PC2 (SI: Table 10). No variables contributed to the model in the best-fit model for PC2 (SI: Table 11). Climate variables contributed ≈10% and ≈20% more than both sex-class and age-class, respectively (SI: Table 12).

## Discussion

This study documents that pigmentation varies with climate across a small geographical scale in a wild lemur population. At Tsinjoarivo, a climate gradient is formed—likely due to the forest being a “high altitude island” and cross-cutting two Köppen climate zones. This unique combination creates a colder and less rainy climate at the highest altitude, which is also the most fragmented habitat. Respectively, both whole body pelage color (Table 1) and facial complexity (Table 2) were associated with climate variables. Specifically, darker individuals with less complex faces also occur most often in this colder, less rainy, more fragmented western forest (Figure 5, Figure 6). However, it is not easy to arrive at any firm conclusions for the mechanism behind the observed pattern without genetic evidence. There are, though, three potential hypotheses that may explain the pattern: (1) isolation by distance, (2) an environmental gradient, or (3) local adaptation. We flesh out intricacies of all three below and set up predictions that would support each hypothesis based on work in other taxa.

The most parsimonious explanation for the pattern in our data is a scenario of isolation by distance (Kimura et al., 1993; Slatkin et al. 1993). First, relatedness may be a confounding factor across our analyses because individuals living in closer proximity are also more likely to be related to each other. This may explain why individuals of similar phenotype cluster together in geographic space (Figure 5). Although we do not have enough data to estimate heritability, from anecdotal observations, we know that offspring can exhibit distinct coloration from their mothers (i.e., fully melanic offspring born to tri-colored mothers). Relatedness aside, forest fragmentation may impede gene flow out of the fragmented forest and create substructure within the population. Thus, individuals in the fragmented forest could also be darker due to inbreeding and/or low genetic diversity. Rare (often melanic) pigmentation phenotypes can arise in other taxa due to increased inbreeding (Larison et al., 2021) or population bottlenecks (Sagar et al., 2021). Although previous work indicates identical mitochondrial haplotypes in groups sampled across Tsinjoarivo, mitochondrial DNA may not always accurately estimate relatedness between individuals or populations (Guschanski et al., 2013; Mayor et al., 2004; Meyer et al., 2016). However, testing the hypothesis of isolation by distance would require genomic evidence we do not yet have. Results supporting this hypothesis may show evidence of restricted gene flow between distinct groups and/or population structure across non-coding and non-outlier loci. An argument against this hypothesis is that there is evidence that sifakas are resilient to fragmentation, as they do not often show decreased gene flow or population structure across fragmented habitats (Quéméré et al., 2010; Salmona et al., 2015; Sgarlata et al., 2016). Sifakas also exhibit high levels of standing genetic variation—the highest observed among primates—despite experiencing a bottleneck ≈100,000 years ago (Guevara et al., 2021). High levels of genome-wide heterozygosity, such as those seen in sifakas, can mean that there is sufficient genetic variation immediately available for selection to act upon when environmental conditions change— thereby increasing the likelihood of rapid adaptive evolution (Barrett & Schluter, 2008).

Thus, this phenotypic pattern may be indicative of selection acting across an environmental gradient. There are two main climate gradients in Madagascar: a North-South gradient in temperature and a West-East gradient in precipitation. Additionally, temperatures decrease as altitudes increase towards the central plateau (Irwin, 2006; Kamilar & Muldoon, 2010). Temperature shows the most significant effect in our models (Table 1, Table 2). Specifically, we found that darker torso and limb color was associated with colder forests with less precipitation (Figure 5). This result supports the thermal melanism hypothesis (i.e., Bogert’s rule) that argues darker fur aids with thermoregulation in colder forests because dark colors may derive more heat from sun rays (Bogert, 1949). Although Bogert’s rule is most commonly used to explain coloration patterns in ectotherms (Bishop et al., 2016; Xing et al., 2016), recent findings extend the rule to birds and mammals (Delhey, 2018; Tapanes et al., 2021). Specifically, Bogert’s rule explains patterns of pelage color variation at a genus-wide level across *Propithecus*, which may implicate thermoregulation as a central selective force in the genus (Tapanes et al., 2021). Tsinjoarivo sifakas, unsurprisingly then, live at the highest altitude and southernmost end of their range. To the North of the diademed sifaka range is the brighter/whiter silky sifaka (*P. candidus*) and to the South is the darker and more melanic Milne-Edwards’s sifaka (*P. edwardsi*) (Mittermeier et al., 2010). Yet evidence for this hypothesis would need to demonstrate that environmental factors, not geographic distance, explain a significant proportion of sifaka genetic variation (McCairns & Bernatchez, 2008).

Another alternative is that, high levels of standing genetic variation (as in sifakas) may increase the efficacy of selection, driving local adaptation within specific populations driven by other selective forces (i.e., not thermoregulation) (Woolridge et al., 2021). A pattern of uniquely locally adapted pelage across Tsinjoarivo is unlikely but would also explain the cluster of darker individuals (as well as less pelage complexity) in colder more fragmented forest. To explain the latter, warm wet regions are assumed to be associated with denser forest habitats—thus, more complex faces in the eastern denser forest may aid with conspecific signal visibility (and potentially inbreeding avoidance (Allen et al., 2014)). However, primate faces exhibit high degrees of phylogenetic signal, thus genetic drift could be driving facial pattern complexity in our data (Bell et al., 2021; Rakotonirina et al., 2017). Other pelage phenotypes (such as degree of redness) may be more salient for signaling, especially given this population of sifakas is polymorphic trichromatic (Jacobs et al., 2017; Tapanes et al., 2021). Nevertheless, under the assumption of high gene flow high across the Tsinjoarivo landscape (as in other sifaka populations), adaptation would have to be strong enough to overcome migration (i.e., selection-migration dynamics) (Haldane, 1930). The mutations underlying phenotypic divergence in this scenario would thus have to be few in number, of large effect, and/or in close proximity (Yeaman & Whitlock, 2011). Thus, genomic evidence for local adaptation within this sifaka population would implicate either one outlier loci of large effect (i.e., *MC1R*, *KITLG*) or many outlier loci of small effect which are gnomically clustered (Pfeifer et al., 2018). All three potential hypothesis should be explored fully with whole exome and/or whole genome data.

However, regardless of the mechanism maintaining phenotypic divergence in this sifaka population, these phenotypic patterns may carry significant implications for conservation. If isolation by distance produces the phenotypic pattern detected here, it would mean that there is limited gene flow between sifakas in fragmented forest and those in continuous forest. Generally, fragmented forest in Tsinjoarivo is low quality habitat and sifakas living there exhibit physiological signs of stress (i.e., low body mass, low white blood cell counts) (Irwin et al., 2008a; 2008b; 2010; 2019). Given predation rates by fosa are higher in fragmented forest (Irwin et al., 2009), isolation may signal the eventual demise of groups living in the fragments. Alternatively, if local adaptation or a larger environmental gradient is responsible for the current pattern, it may foretell future maladaptation and potentially local extinction for the population—because loss of habitat where a population has been uniquely adapted to thrive may drive environmental mismatch. Such a mismatch can have consequences, such as increased predation on maladapted phenotypes (e.g., a mismatch between pelage and background color). Either scenario thus poses a significant risk to the already Critically Endangered diademed sifaka and the unprotected forest of Tsinjoarivo (which harbors high levels of biodiversity) (Goodman et al., 2000; Irwin, 2020). It’s likely that across other populations, hair variation may also serve as a signal of adaptation, environmental mismatch, or restricted gene flow. However, such data is missing in the primatological literature, thus, it remains unknown if the patterns detected across Tsinjoarivo sifakas are exceptional or typical.

## Acknowledgements

We thank the Government of Madagascar (Ministry of Environment and Sustainable Development) for permission to carry out the project, and SADABE and the University of Antananarivo Mention Anthropobiologie et Développement Durable for facilitation and partnership. We thank Ian Harryman for assistance with scoring sifaka pelage, and Jean-Luc Raharison, Bruno Ramorasata, Narcisse Rahajanirina and Tsinjoarivo research technicians for animal capture logistic assistance, and Karine Lalaina Mahefarisoa, Mamy Navalona Andriamihajarivo and Pierre Michel Ralaivelo for veterinary services. The project was funded by National Science Foundation (BCS #1354997, and BCS #1636360), International Primatological Society, Explorer’s Club, The George Washington University, Northern Illinois University, National Geographic Society, Eppley Foundation for Research, IdeaWild, Rasmussen Family Foundation, St. Louis Zoo, Primate Conservation, Inc., Primate Action Fund, Margot Marsh Biodiversity Foundation, and Stony Brook University. We thank Timothy Webster and Andrew Zamora for feedback on earlier drafts of this manuscript, as well as anonymous reviewers and an editor that provided useful feedback for us to effectively revise a previous version of this manuscript.

