## Supplementary material for "Melanism in a wild sifaka population: Darker where cold and fragmented": SI:

**Table 1** WorldClim bioclimatic variables used in our ENMs.

| Layer Code | Bioclimatic Variable |
|---|---|
| BIO 03 | Isothermality |
| BIO 04 | Temperature Seasonality |
| BIO 07 | Temperature Annual Range |
| BIO 11 | Mean Temperature of Coldest Quarter |
| BIO 16 | Precipitation of Wettest Quarter |
| BIO 19 | Precipitation of Coldest Quarter |

For more detailed descriptions, visit: http://worldclim.org/bioclim.

**Table 2** Bioclimatic variables contributing to ENMs for Tsinjoarivo.

| **Model** | **Variable** | **Percent Contribution** | **Permutation Importance** |
|---|---|---|---|
| Fragmented sites | Bio11 | 77.7 | 12.9 |
| | Bio07 | 15.8 | 33.5 |
| | Bio16 | 2.9 | **38.3** |
| | Bio04 | 2.1 | 0 |
| | Bio19 | 1.4 | 15.4 |
| | Bio03 | 0 | 0 |
| Continuous sites | Bio11 | 86.5 | **55.9** |
| | Bio07 | 6.8 | 32.4 |
| | Bio16 | 5.5 | 3 |
| | Bio04 | 1.2 | 8.7 |
| | Bio19 | 0 | 0 |
| | Bio03 | 0 | 0 |

**Table 3** Distribution of sampling across regions, age-class, and sex-class in large Tsinjoarivo sample, for larger semi objective scoring methodology (Figure 3).

| Age-Class: Sex Class | Region | | | |
|---|---|---|---|---|
| | **VATA** | **ANKA** | **MAHA** | **All** |
| **AD: F** | 5 | 4 | 6 | 15 |
| **AD: M** | 5 | 1 | 10 | 16 |
| **SA: F** | 6 | 4 | 7 | 17 |
| **SA: M** | 8 | 5 | 8 | 21 |
| **JUV: F** | 1 | 3 | 3 | 7 |
| **JUV: M** | 4 | 2 | 5 | 11 |
| **All** | 29 | 19 | 39 | 87 |

Key: Adults (AD), sub-adults (SA), and juveniles (JUV) numbers, as well as Vatateza (VATA), Ankadivory (ANKA), and Mahatsinjo (MAHA).

**Table 4** Pelage variables scoring definitions for coloration variables, countershade, complexity, and contrast.

| Pelage Variable | Scoring Definition |
|---|---|
| Head pigmentation | Mean: (A+B+C+D+E+F)/6 |
| Forelimb pigmentation | Mean: (G+H)/2 |
| Hindlimb pigmentation | Mean: (L+M)/2 |
| Tail pigmentation | Mean: (O + P)/2 |
| Dorsal torso pigmentation | Mean: (I+K)/2 |
| Ventral torso pigmentation | Mean: (Q+R)/2 |
| Countershade | Difference between dorsal torso pigmentation and ventral torso pigmentation |
| Head complexity | # of distinct shades between A and F |
| Anterior body complexity | # of distinct shades between G and P |
| Ventral complexity | # of distinct shades between Q and R |
| Head contrast | Difference between highest and lowest score between A and F |
| Anterior body contrast | Difference between highest and lowest score between G and P |
| Ventral contrast | Difference between highest and lowest score between Q and R |

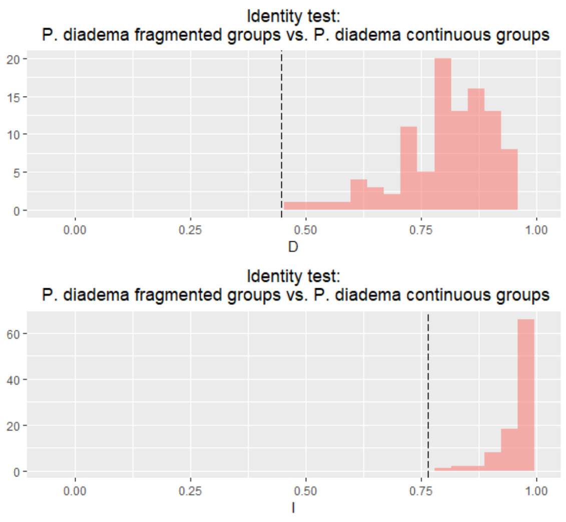


**Figure 1** Distribution of *Schoener's D* and *Hellinger's I* for 99 pseudo-habitat pairs.

**Table 5** Full results of identity test.

| Test | P-value | Niche Overlap Values |
|---|---|---|
| Schoener's D | 0.01 | 0.4915 |
| Hellinger's I | 0.01 | 0.8148 |

**Table 6** Full results from the binomial test of omission.

| Data | Fold | Omission | P-value | N train | N test |
|---|---|---|---|---|---|
| Cont. | 1 | 0.50 | 8.98E+03 | 3 | 2 |
| Cont. | 2 | 0.50 | 3.97E+05 | 3 | 2 |
| Cont. | 3 | 0.00 | 3.84E+05 | 3 | 2 |
| Cont. | 4 | 0.00 | 4.09E+05 | 3 | 2 |
| Frag | 1 | 0.67 | 2.68E+02 | 4 | 3 |
| Frag | 2 | 0.33 | 2.41E+04 | 4 | 3 |
| Frag | 3 | 0.33 | 2.26E+04 | 4 | 3 |
| Frag | 4 | 0.33 | 2.41E+04 | 4 | 3 |

**Table 7** Best models predicting pelage coloration for PC1 using Akaike's Information Criterion with cor
small sample sizes (AICc)

| **Model** | **Independent Variables** | | | | **df** | **AICc** | **Δ AICc** | **AICc weight** |
|---|---|---|---|---|---|---|---|---|
| | *AgeClass* | *BIO11* | *BIO16* | *Sex* | | | | |
| PC1-a | -0.307 | 18.110 | -0.331 | | 5 | 308.5 | 0 | 0.392 |
| PC1-b | | 18.510 | -0.337 | | 4 | 308.7 | 0.16 | 0.361 |
| PC1-c | -0.311 | 18.020 | -0.330 | + | 6 | 310.8 | 2.24 | 0.128 |
| PC1-d | | 18.460 | -0.337 | + | 5 | 310.9 | 2.39 | 0.118 |

Key: AgeClass = adult (3), sub-adult (2), or juvenile (1); Sex = male or female. A '+' sign indicates that t
variable is included in the model, and numerical values indicate the coefficient and that a variable contrib
model. All models within two AICc values from the best model are considered equivalently good models
represents the 'log-likelihood ratio chi$^2$'.

**Table 8** Sum of AICc weights of predictor variables for models predicting pelage coloration of PC1 in pigmentation PCA.

| **Variables:** | *BIO16* | *BIO11* | *AgeClass* | *SexClass* |
|---|---|---|---|---|
| **Sum of Weights** | 1.00 | 1.00 | 0.52 | 0.25 |
| **N containing models** | 8 | 8 | 8 | 8 |

**Table 9** Best models predicting pelage color patterning across PC1 using Akaike's Information Criterion corrections for small sample sizes (AICc)

| **Model** | **Independent Variables** | | | | **df** | **AICc** | **Δ AICc** | **AICc weight** | |
|---|---|---|---|---|---|---|---|---|---|
| | *AgeClass* | *BIO11* | *BIO16* | *Sex* | | | | | |
| PC1-a | | -7.879 | 0.131 | | 4 | 294.8 | 0 | 0.263 | - |
| PC1-b | | | | | 2 | 296.6 | 1.84 | 0.105 | - |
| PC1-c | 0.113 | -7.734 | 0.129 | | 5 | 296.6 | 1.87 | 0.103 | - |

Key: Regions = Mahatsinjo, Ankadivory, Vatateza; AgeClass = adult, aub-adult, or juvenile; Sex = male '+' sign indicates that the categorical variable is included in the model, and numerical values indicate the and that a variable contributed to the model. All models within two AICc values from the best model are equivalently good models. Loglik represents the 'log-likelihood ratio chi$^2$'.

**Table 10** Sum of AICc weights of predictor variables for models predicting pelage variation of PC1 in patterning PCA.

| **Variables:** | *BIO11* | *BIO16* | *AgeClass* | *SexClass* |
|---|---|---|---|---|
| **Sum of Weights** | 0.66 | 0.62 | 0.30 | 0.25 |
| **N containing models** | 8 | 8 | 8 | 8 |

**Table 11** Best models predicting pelage color patterning across PC2 using Akaike's Information Criterion with corrections for small sample sizes (AICc).

| **Model** | **Independent Variables** | | | | **df** | **AICc** | **Δ AICc** | **AICc weight** | **LogLik** |
|---|---|---|---|---|---|---|---|---|---|
| | *AgeClass* | *BIO11* | *BIO16* | *Sex* | | | | | |
| PC2-a | | | | | 2 | 275.6 | 0 | 0.133 | -135.704 |
| PC2-b | | 6.457 | -0.115 | | 4 | 275.7 | 0.17 | 0.122 | -133.616 |
| PC2-c | -0.217 | | | | 3 | 276.0 | 0.46 | 0.105 | -134.863 |
| PC2-d | | | | + | 3 | 276.4 | 0.85 | 0.087 | -135.056 |
| PC2-e | -0.200 | 6.203 | -0.111 | | 5 | 276.5 | 0.94 | 0.083 | -132.875 |
| PC2-f | -0.231 | | | + | 4 | 276.7 | 1.12 | 0.076 | -134.092 |
| PC2-g | | 6.183 | -0.110 | + | 5 | 277.0 | 1.41 | 0.066 | -133.109 |
| PC2-h | -0.214 | 5.883 | -0.105 | + | 6 | 277.6 | 2.01 | 0.049 | -132.253 |
| PC2-i | | | -0.002 | | 3 | 277.6 | 2.06 | 0.048 | -135.697 |

Key: Regions = Mahatsinjo, Ankadivory, Vatateza; AgeClass = adult, aub-adult, or juvenile; Sex = male or female. A '+' sign indicates that the categorical variable is included in the model, and numerical values indicate the coefficient and that a variable contributed to the model. All models within two AICc values from the best model are considered equivalently good models. Loglik represents the 'log-likelihood ratio $chi^2$'.

**Table 12** Sum of AICc weights of predictor variables for models predicting pelage variation of PC2 in patterning PCA.

| **Variables:** | *BIO16* | *BIO11* | *AgeClass* | *SexClass* |
|---|---|---|---|---|
| **Sum of Weights** | 0.46 | 0.46 | 0.44 | 0.39 |
| **N containing models** | 8 | 8 | 8 | 8 |
